# Structure and Post-Translational Modification of the Prostaglandin Transporter

**DOI:** 10.1101/2024.12.31.630934

**Authors:** Peixuan Yu, Melanie A. Orlando, Benjamin J. Orlando

**Affiliations:** Dept. of Biochemistry and Molecular Biology, Michigan State University, USA

**Keywords:** PGT, OATP2A1, disulfide, glycosylation, prostaglandin

## Abstract

The prostaglandin transporter (PGT) is a member of the Organic Anion Transporting Polypeptide (OATP) family of membrane transporters. PGT mediates the uptake of prostaglandins from the extracellular environment to enable intracellular enzymatic degradation and termination of signaling. In addition to importing prostaglandins, PGT is also an essential core component of the Maxi-Cl channel, which facilitates cellular release of ATP and other small organic anions. Despite progress on understanding the (patho)physiological roles of PGT, and development of small molecules to inhibit this transporter, molecular details of the overall structure and transport mechanism remain elusive. Here we determined the cryo-EM structure of human PGT, which demonstrates an overall topology consistent with other OATPs despite possessing a dual transporter/channel functionality. We additionally investigated the role of eight potential disulfide bonds found in the extracellular loops of PGT and paralogous transporters. Through biochemical and functional characterization we demonstrate that six intra-loop disulfide bonds (C420-C511, C450-C470, C492-C474, C459-C507, C444-C494, C580-C594) are essential for proper N-glycosylation, plasma membrane trafficking, and prostaglandin import activity of PGT. In contrast, two inter-loop disulfides (C155-C587 and C143-C448) were found to restrict maximal prostaglandin uptake, suggesting a possible regulatory role in modulating PGT activity. In total, our studies provide a fresh molecular perspective on the structure, post-translational modification, and overall function of PGT.

**Significance Statement:** The prostaglandin transporter (PGT) is essential for cellular uptake of prostaglandins and serves as a core component of the Maxi-Cl channel. Using cryo-EM we resolved the structure of human PGT, revealing a similar overall topology as other OATP transporters. Extensive site-directed mutagenesis and functional assays identified eight disulfide bonds in PGT’s extracellular domain as key regulators of glycosylation, trafficking, and transport activity. These findings provide a structural basis for PGT function and lay a foundation to further explore substrate recognition and inhibitor design for this biologically significant transporter.

## Introduction

Prostaglandins (including PGE_2_, PGF_2α_, PGD_2_, PGI_2_, and TxA_2_) are bioactive oxygenated fatty acids that play crucial roles as signaling molecules throughout the human body. Prostaglandin signaling mediates a wide range of processes such as pain and inflammation^1^, ovulation^2,3^, blood pressure regulation^4^, and cancer progression^5^. Prostaglandin homeostasis is maintained through a dynamic interplay of biosynthesis, transporter mediated cellular export and import, and intracellular enzymatic degradation. After being synthesized intracellularly by the cyclooxygenase enzymes and tissue specific synthases, prostaglandins are exported from the cell via active transporters such as multiple drug resistance-associated protein 4 (MRP4)^6–8^. Once exported to the extracellular environment, prostaglandins initiate diverse signaling pathways by binding to a repertoire of different G-protein coupled receptors^9^. To terminate signaling, prostaglandins are re-imported by the prostaglandin transporter (PGT), and enzymatically inactivated by 15-hydroxyprostaglandin dehydrogenase (15-PGDH)^10^(**Suppl. Fig. S1A**). In humans it has been shown that PGT is ubiquitously expressed and is the primary transporter mediating prostaglandin import^11,12^.

Accumulating evidence suggests that PGT is involved in many mammalian physiological processes. For example, *Slco2a1* (gene encoding PGT)-knockout mouse models show defects in prostaglandin entry to tissues such as the lungs, gastrointestinal tract^13^, and placenta^14^. Additionally, *Slco2a1* knockout mice show failure of the ductus arteriosus to close within 48 hours post-birth, resulting in lethal patent ductus arteriosus (PDA) that can be overcome through indomethacin treatment^15^. Whole-exome analyses also suggest that recessive inheritance of *SLCO2A1* mutations are associated with primary hypertrophic osteoarthropathy (PHO)^16^ and chronic enteropathy associated with *SLCO2A1* (CEAS)^17^. Moreover, PGT is considered a potential pharmacological target for treating diabetic foot ulcers^18^, antipyresis, and non-hormonal contraception^2^. Small molecule inhibitors of PGT^19^ have been developed and show promise in accelerating wound healing in non-diabetic and diabetic rats^18^. Beyond its role as a transporter, PGT has also been identified as a core component of the Maxi-Cl channel when the transporter is complexed with AnnexinA2-p11^20,21^. In this capacity, PGT can facilitate the cellular release of ATP and other small organic anions. While the physiological roles of PGT have been well established, the mechanism by which the transporter mediates prostaglandin import and Maxi-Cl channel activity remain unclear.

PGT is a member of the Organic Anion Transporter Polypeptide (OATP) family of transporters^22^. All 11 human OATP transporters are predicted to contain a 12-transmembrane spanning helix architecture, with a Kazal domain in extracellular loop 3 (ECL3) that extends between transmembrane helix 9 and 10 (**Fig. 1A&B, Suppl. Fig S2**). A notable feature of the extracellular domains across the OATP family is the presence of 16 conserved cysteine residues, which form eight pairs of disulfide bonds as demonstrated by previous free cysteine labeling assays^23^ and the recent cryo-EM structures of OATP1B1 and OATP1B3^24,25^. Several missense mutations in these extracellular cysteines (C420F^26^, C459R^27^, C492Y^28^, C594Y^29^) have been identified in patients with CEAS and PHO. However, the mechanisms by which these cysteine missense mutations lead to PGT dysfunction and resultant PHO and CEAS remains unclear. In other OATPs and similar transporters, disulfide bonds in extracellular domains have been shown to be essential for proper subcellular localization and/or transport function. For example, five pairs of disulfide bonds in the longest ECL of OATP2B1^23^ are important for the proper targeting of this transporter to the plasma membrane. Similarly, cysteine to alanine substitutions at four locations in the extracellular region of murine OAT1^30^ impairs trafficking to the cell surface and leads to almost complete loss of function. Moreover, PGT undergoes extensive glycosylation in ECL3, which is critical for proper membrane localization and function of the transporter^31^. Together these studies suggest that PGT undergoes a series of complex post-translational modifications that have significant effects on subcellular localization and transporter function.

**Figure 1.**
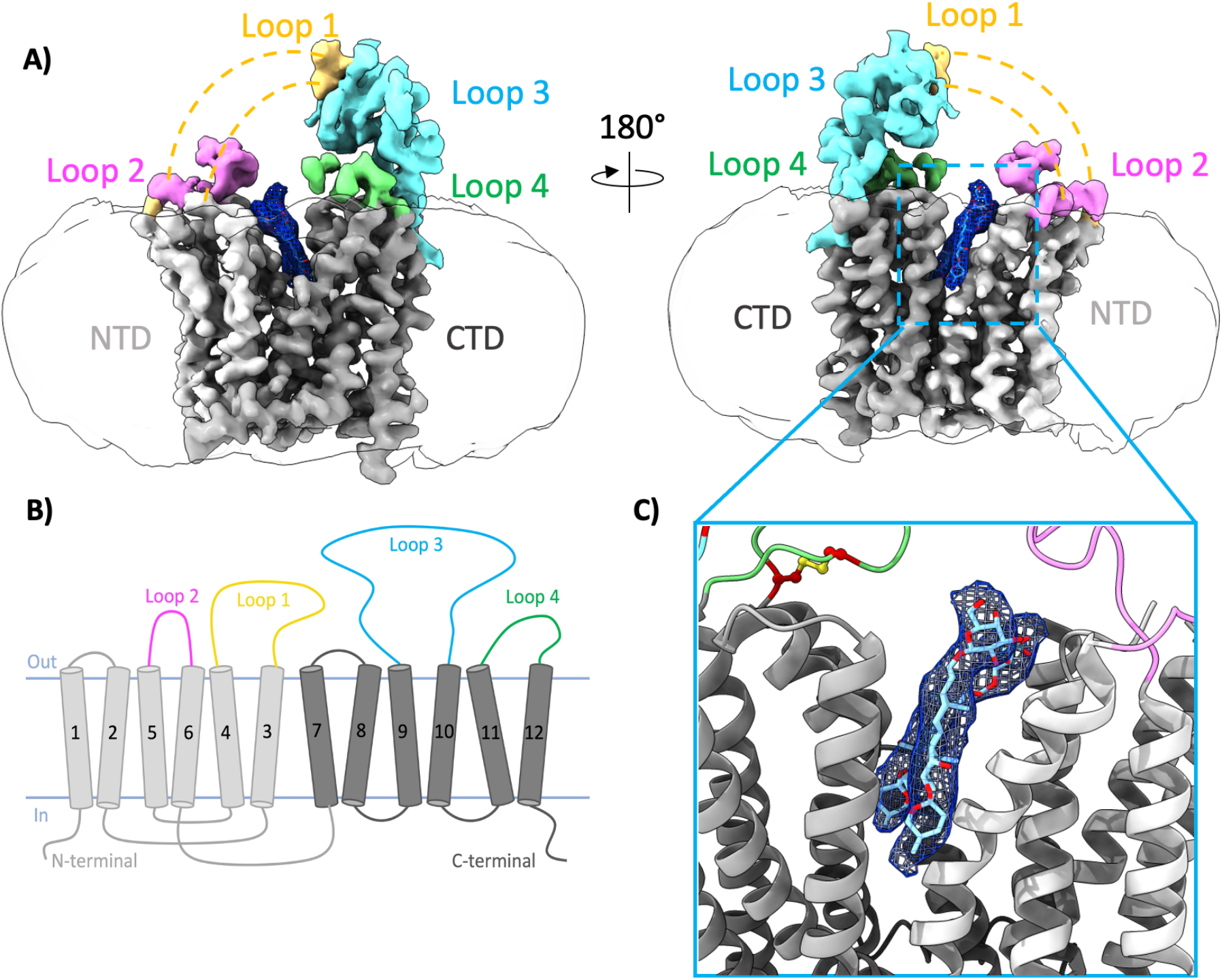
The overall architecture of PGT. **A)** Two rotated views of the cryo-EM map of PGT. The N-terminal (NTD) and C-terminal domain (CTD) are colored in light grey and dark grey, respectively. Extracellular loop 1, 2, 3, 4 are colored in yellow, pink, cyan and green, respectively. The detergent micelle is shown in light transparent white, and digitonin-like density is shown in dark blue. Dashed lines show the flexible regions in our structures. **B)** The membrane topology of PGT. **C)** A detergent-like density at the transport pocket between NTD and CTD. Digitonin (light cyan) is fit into this density.

In this study we present the cryo-EM structure of human PGT in an outward-facing state. The structure reveals that PGT adopts the same topology as other OATPs despite the ability to function as a bimodal transporter/channel. We further characterized the role of individual disulfide bonds in the ECD through analysis of glycosylation, membrane trafficking, and prostaglandin transport. Together, the cryo-EM structure and associated functional analyses provide key insight into the overall structure and topology of PGT, potential ligand binding pockets, and the role of extracellular disulfide bonds in facilitating proper PGT localization and function. These results shed light on the impact of cysteine missense mutations in patients with PHO and CEAS, and also provide a structural platform upon which to begin planning rational improvement of PGT selective small molecule transport inhibitors.

## Results

### The overall architecture of PGT

Recent cryo-EM analysis of the human liver transporters OATP1B1 and OATP1B3 have provided key insight into the overall structure and conformational cycle of OATP transporters^24,25^. Based on sequence similarity (**Suppl. Fig. S2**) and publicly available structural predictions (AlphaFold Protein Structure Database), PGT and the other 10 human OATPs are likely to adopt a similar overall topology to that observed for OATP1B1 and OATP1B3. To confirm this prediction, we pursued a cryo-EM structure of human PGT. To obtain transporter preparations of sufficient quality and quantity for structural studies, a HEK293-F stable cell line expressing N-terminal twin-strep tagged human PGT was constructed (see methods). PGT was isolated from this cell line using a Streptactin-XT affinity column with on-column detergent exchange and final size-exclusion chromatography in digitonin. This procedure generated highly pure and monodisperse samples amenable for structural studies **(Suppl. Fig. S1B&C)**, and the resulting cryo-EM images of detergent solubilized PGT show a clear distribution of purified particles (**Suppl. Fig. S1D&E**).

The cryo-EM map of human PGT (**Fig. 1A-C**) was resolved at an intermediate overall resolution of 4.3Å using single-particle cryo-EM **(Suppl. Fig. S3-4)**. The extracellular loops of PGT and the extracellular apex of the N-terminal bundle of transmembrane helices (TM_1-6_) are not as clearly resolved as the rest of the transporter (**Fig. 1A&B, Suppl. Fig. S4C&D**). Despite the intermediate overall resolution, the cryo-EM map is of sufficient quality to visualize most sidechains in the transmembrane helices, allowing for unambiguous docking and refinement of an atomic model generated with AlphaFold^35^. Comparison of the final atomic model generated from fitting of the cryo-EM data with that generated from the Alphafold prediction alone (**Suppl. Fig. S5A**) reveals striking similarity (RMSD ∼1.6Å over 465 C_α_ atoms), indicating a high degree of accuracy in the Alphafold model. The largest difference between experimentally derived and predicted models is in the degree of opening between the two transmembrane regions, with the cryo-EM derived model showing a slightly larger opening between N-terminal and C-terminal transmembrane bundles **(Suppl. Fig. S5C)**.

In our cryo-EM structure, PGT adopts an outward-facing conformation similar to that observed for human OATP1B1 **(Suppl. Fig. S5A)**. Our cryo-EM structure thus confirms that PGT, an OATP with both transporter and channel-like activity, adopts the classic major facilitator superfamily (MFS) fold with 12 transmembrane helices (TMs) organized into distinct N-terminal (NTD) and C-terminal (CTD) domains related by 2-fold pseudosymmetry **(Fig. 1A&B)**. Prior to freezing grids for cryo-EM analysis we incubated purified PGT with the transport substrate PGE_2_. Although density for PGE_2_ was not observed in the cryo-EM map, flat detergent like densities were found in the hydrophobic cleft between the NTD and CTD **(Fig. 1C)**. Based on their shape and location within the expected membrane plane, these densities were assigned as digitonin detergent molecules. The positioning of these detergent molecules between the NTD and CTD transmembrane bundles likely facilitated structure determination by sterically blocking transition of PGT towards an inward facing or occluded state (**Suppl. Fig. S5A&B**). Moreover, the digitonin is positioned directly above and just outside the substrate binding pocket identified for bilirubin, semiprivir, and dichlorofluorescein binding in OATP1B1^24^**(Suppl. Fig. S6)**.

Although we were not able to visualize the transport substrate PGE_2_ bound in our cryo-EM map, previous biochemical analyses have delineated the approximate substrate binding site. Cysteine scanning mutagenesis combined with covalent cysteine modification assays have demonstrated that PGE_2_ likely binds in a pocket near the middle of TM_10_ by residues S526-H533 ^45^**(Suppl. Fig. S7A)**. Charged residues R561 and K614 in the 11^th^ and 12^th^ TM helices respectively, have been proposed as critical residues for substrate binding and transport^46^. In an approach analogous to that recently utilized for OATP1B1 and OATP1B3^24,25^, we utilized molecular docking to gain insight into the possible binding poses of PGE_2_ within the PGT transport pathway. Autodock Vina^41^ was utilized to dock PGE_2_ into the cryo-EM structure of PGT after removal of the digitonin molecules, and the six of the amino acids described above were designated as flexible residues **(Suppl. Fig. S7B)**. The top ranked PGE_2_ binding poses show the negatively charged carboxyl group of the ligand positioned away from Lys614 in TM_12_ and closer to Arg561 in TM_11_ **(Suppl. Fig. S7C-D)**. This suggests that Arg561 may be critical in facilitating PGE_2_ binding, while Lys614 may facilitate binding of smaller molecules such as lactate^47^, which have been identified as counter substrates to drive PGE_2_ uptake. Additionally, most of the observed binding poses agree with previous cysteine scanning studies that identified S526, A529, C530, and H533 as critical residues that line the PGE_2_ binding pocket **(Suppl. Fig. S7C&D)**^45^. These PGE_2_ binding poses are also consistent with the binding sites identified for bilirubin and semiprevir in OATP1B1^24^ **(Suppl. Fig. S7F&G)**. However, in one PGE_2_ binding pose identified from molecular docking, the bound PGE_2_ is slightly offset from the central pocket and positioned more towards the NTD (**Suppl. Fig. S7E&H)**. While binding of PGE_2_ in this position is less supportive of previous biochemical assays that demonstrate residues S526-H533 in TM_10_ line the binding pocket, we cannot exclude that this may be a transient PGE_2_ binding position. Further structural analysis will be required to determine precisely how prostaglandins and other small molecules are recognized by PGT, and to elucidate the precise roles of charged residues R561 and K614 in mediating this process. During preparation of this manuscript a cryo-EM map (EMDB 37234) and atomic coordinates (PDB 8KGW) for PGT bound to PGE_2_ were publicly released. While a full description of this map and coordinates awaits formal publication, the binding location of PGE_2_ observed in this newly released model is highly similar to one of the poses identified from molecular docking (**Suppl. Fig. S7D**).

In both our cryo-EM map and those of OATP1B1 and OATP1B3^24,25^, loops comprising the extracellular domain (ECD) are significantly more disordered and lower resolution than the TM region (**Fig. 1A-C, and Suppl. Fig. S4C&D**). The ECD of PGT is composed of four extracellular loops (**Fig. 1B and 2A**), with the longest loop, loop-3, containing a glycosylated serine protease-like Kazal domain of largely unknown function. In the cryo-EM map, the Kazal domain of loop-3 is positioned directly over the CTD transmembrane bundle (**Fig. 1A and 2A**) in a configuration that is analogous to that seen with OATP1B1 in an outward facing state^24^ (**Suppl. Fig. S5A**). The limited resolution and fragmented densities of the ECD in the PGT cryo-EM map prevent precise atomic model building in this region, with loop-1 being almost completely disordered (**Fig. 1A&B**). However, several factors support the utility of an outward-facing AlphaFold model of PGT for mechanistic insights. These include PGT’s homology to other OATPs of known structure, AlphaFold’s striking accuracy in modeling the TM region of PGT, and sequence-specific features discussed below.

### Disulfide bonds in the extracellular domain of PGT

A surprising feature in the ECD of human OATPs is the presence of 16 conserved cysteine residues (**Fig. 2C**). These residues are found throughout the loops of the ECD except for loop-2, which is the only extracellular loop that is completely devoid of cysteines (**Figure 2A-C**). Despite the lower overall confidence (pLDDT score) of the ECD in Alphafold models of PGT and other OATPs (**Suppl. Fig. S5D**), the 16 conserved cysteine residues are invariably positioned within proper distance from one another to support formation of 8 total disulfide bonds (**Fig. 2B**). The predicted disulfide bond pattern is largely supported by those disulfides (C430-C530, C599-C613, C489-C504, C142-C463, C459-C506, C474-C524) that are directly visible in the higher resolution OATP1B1^24^ and OATP1B3^25^ structures. In PGT, five intra-loop disulfide bonds (C474-C492, C459-C507, C444-C494, C450-C470, C420-C511) are predicted in the loop-3 Kazal domain^48^, and one intra-loop disulfide (C580-C594) is predicted to staple the two termini of loop-4 together (**Fig. 2B&C**). In contrast, two other pairs of cysteines are predicted to form inter-loop disulfide bonds between loop-1 and loop-3 (C143-C448) or loop-1 and loop-4 (C155-C587). To further analyze these conserved cysteines and putative disulfide bonds, we undertook systematic site-directed mutagenesis to probe the role of each individual cysteine.

**Figure 2.**
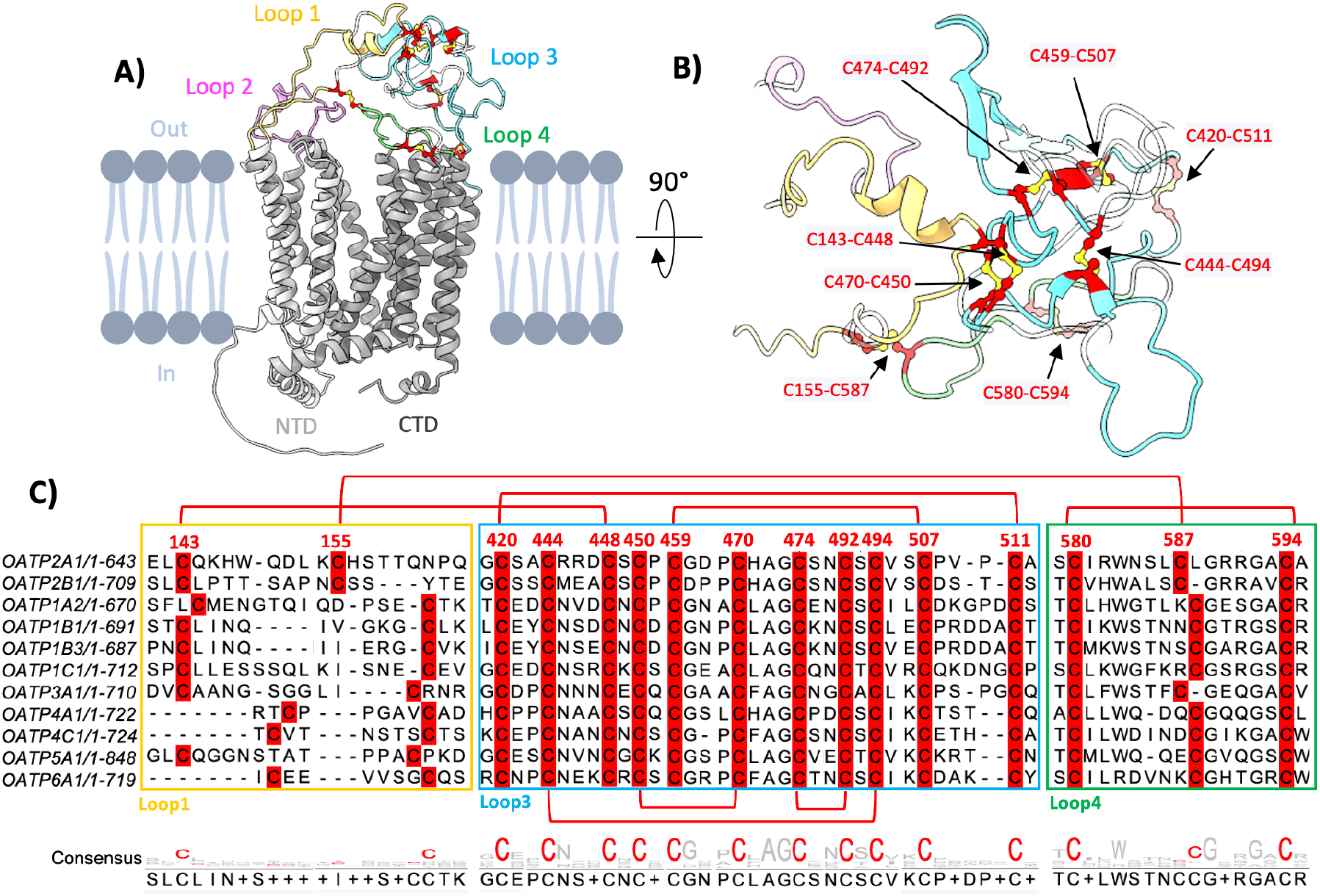
Disulfide bonds in the extracellular domain of PGT. **A)** AlphaFold atomic model of human PGT in an outward facing conformation. The N-terminal (NTD) and C-terminal domain (CTD) are colored in light grey and dark grey, respectively. Extracellular loop 1, 2, 3, 4 are colored in yellow, pink, cyan and green, respectively. **B**) Disulfide pattern in the extracellular domain of the AlphaFold-predicted human PGT model. Eight disulfide bonds are colored in red with sulfur atoms in yellow. **C)** The sequence alignment of OATP transporters in the section containing the 16 cysteine residues. Disulfide bond pairs are indicated by red lines, and sequences of loop 1, 3, and 4 are circled in yellow, cyan and green box, respectively.

We evaluated the whole-cell expression levels of wild-type PGT and 16 individual Cys-to-Ser variants, each transiently transfected into HEK293-F cells. Expression levels were analyzed using anti-HA tag western blotting of whole-cell lysates. All 16 Cys-to-Ser variants showed robust expression in HEK293-F cells (**Fig. 3A-C**). However, notable differences in banding patterns were observed, indicating possible alterations in glycosylation among the variant transporters. To further investigate, we expressed WT PGT in HEK293 GnTI (−) cells, which lack N-acetylglucosaminyltransferase I (GnTI), a Golgi resident enzyme required for synthesizing hybrid and complex N-glycans^49^. In these GnTI (−) cells, transfected WT PGT produced only a single lower band at ∼75 kDa (**Fig. 3A-C**). In contrast, WT PGT expressed in HEK293-F cells exhibited two distinct bands on the western blot: an upper band at ∼100 kDa corresponding to fully glycosylated PGT and a lower band at ∼75 kDa representing partially glycosylated PGT^31^ (**Fig. 3A-C**).

**Figure 3.**
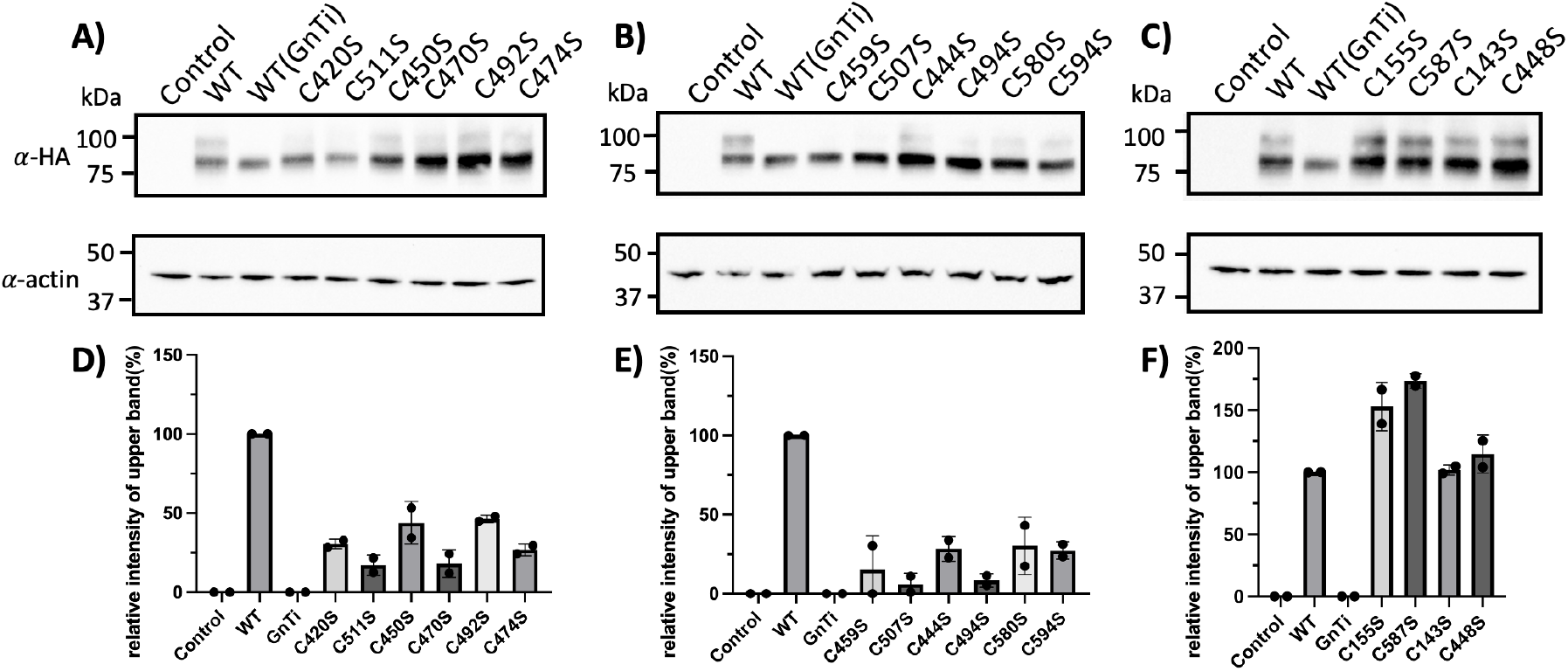
Whole-cell expression of human PGT cysteine variants. **A, B, C)** Immunoblot of whole cell expression levels of WT PGT and cysteine variants. The upper band represents the fully glycosylated PGT, and the lower band represents PGT in a high-mannose glycan form. **D, E, F)** ImageJ analysis of the relative intensity ratio between the upper band and the total intensity of both upper and lower bands. Errors bars represent standard deviation of the mean (SD) across *n* = 2 replicate measurements.

Compared to WT PGT, the C459S and C507S variants did not display any of the mature glycoform (**Fig. 3B&E**), indicating inefficient maturation of glycosylation in the Golgi apparatus for these variants. Ten other cysteine variants (C420S, C511S, C450S, C470S, C492S, C474S, C444S, C494S, C580S, C594S) show greatly reduced quantities of the mature glycoform (**Fig. 3A&B and 3D&E**), suggesting defects in maturation of glycosylation for these variants as well. In contrast, the remaining four Cys-to-Ser variants (C155S, C587S, C143S, C448S) expressed both the immature and mature glycoforms, and displayed a similar or even slightly elevated degree of the mature glycoform compared to WT (**Fig. 3C&F**). In total, these results suggest that removal of intra-loop disulfide bonds between C420-C511, C444-C494, C450-C470, C459-C507, and C464-C492 negatively impacts or even prevents proper maturation of PGT glycans, whereas removal of the inter-loop disulfide bonds between C143-C448 and C155-C587 promoted or had no effect on glycan maturation.

### Surface expression and transport activity of PGT cysteine variants

Previous studies demonstrated that mutating N-glycosylation sites of PGT^31^ and OATP2B1^50^ negatively impacts the ability of these transporters to be trafficked to the plasma membrane. We hypothesized that PGT cysteine variants exhibiting reduced levels of the mature glycoform (**Fig. 3A-F**) may also display impaired trafficking to the plasma membrane. To test this hypothesis a surface biotinylation assay^23^ was used to evaluate the surface expression levels of WT PGT and cysteine variants. The cell-impermeant sulfo-NHS-SS-biotin reacts with primary amines and was used to label WT PGT and its cysteine mutants on the plasma membrane for subsequent pulldown and western blot detection. WT PGT-transfected cells without sulfo-NHS-SS-biotin labeling and empty vector-transfected cells with sulfo-NHS-SS-biotin labeling served as negative controls (**Fig. 4A**). No bands were observed for these negative controls, indicating that the assay does not capture any unlabeled PGT. Therefore, bands observed on the immunoblot represent PGT located on the plasma membrane. For WT PGT expressed in HEK293 GnTi(−) cells, a lower band appears on the immunoblot, indicating that the immature glycoform expressed in GnTi(−) cells can be trafficked to the plasma membrane **(Fig. 4A&B)**. Among the 12 intra-loop cysteine variants that display a reduction or full abolishment of the mature glycoform **(Fig. 3A&B**), 11 variants (C420S, C511S, C470S, C492S, C474S, C459S, C507S, C444S, C494S, C580S, C594S) show very faint bands in the surface biotinylation assay (**Fig. 4A&B**), indicating that these variants cause significant defects in proper plasma membrane localization of PGT. The C450S variant displays a slightly stronger band than the other 11 cysteine variants in the surface biotinylation assay, indicating a minimal amount of expression at the plasma membrane (**Fig. 4A**). In contrast, the four inter-loop cysteine variants (C155S, C587S, C143S, and C448S) that displayed normal levels of the mature glycoform (**Fig. 3C&F**) also display robust labeling in the surface biotinylation assay (**Fig. 4C**), demonstrating that these variants are both glycosylated and properly trafficked to the plasma membrane. In total, these results demonstrate that six out of the eight putative disulfide bonds (C420-C511, C450-C470, C492-C474, C459-C507, C444-C494, C580-C594) in the ECD of PGT are required for proper maturation of glycosylation and trafficking to the plasma membrane.

**Figure 4.**
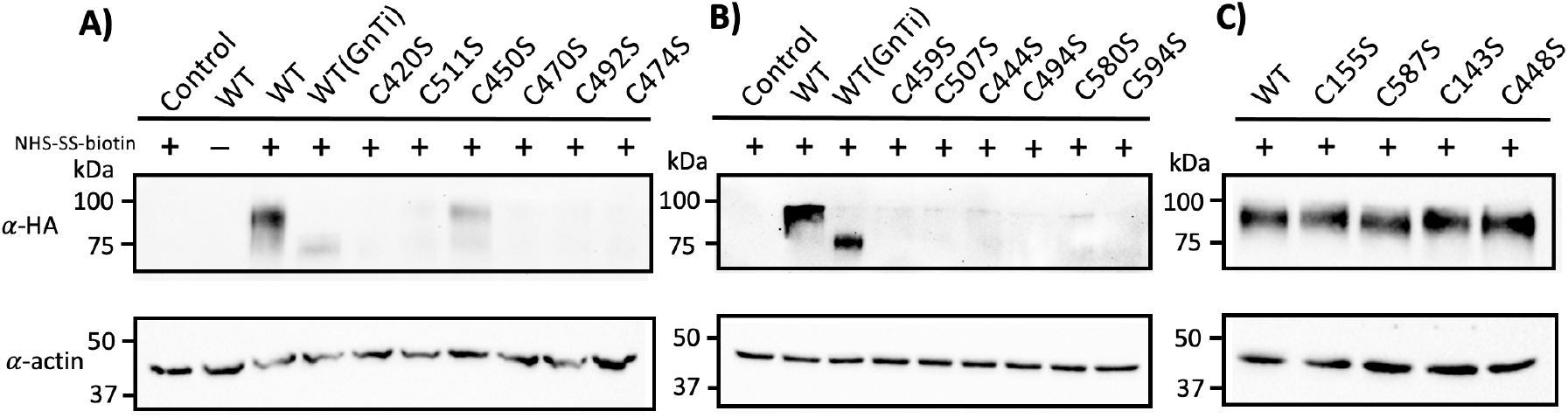
Surface expression of PGT cysteine variants using sulfo-NHS-SS-biotin. **A)** Immunoblot of sulfo-NHS-SS-biotin labeled WT PGT and cysteine variants C420S, C511S, C450S, C470S, C492S, C474S, probed with anti-HA antibody. Anti-actin immunoblotting was used as a loading control. **B)** Anti-HA immunoblot of sulfo-NHS-SS-biotin labeled PGT WT and cysteine variants C459S, C507S, C444S, C494S, C580S, C594S. Anti-actin immunoblotting was used as a loading control. **C)** Immunoblot of sulfo-NHS-SS-biotin labeled WT PGT and cysteine variants C155S, C587S, C143S, C448S.

### PGE_2_ uptake activity of PGT cysteine variants

From the surface biotinylation results described above, it is reasonable to assume that cysteine variants displaying reduced or abolished plasma membrane localization of PGT should also display reduced or abolished transport of prostaglandins. To test this hypothesis we performed mass-spectrometry based prostaglandin uptake assays using PGE_2_-d_4_ as the substrate and PGE_2_-d_9_ as an internal quantification standard^43^. In this assay the intra-loop cysteine variants (C420S, C511S, C470S, C492S, C474S, C459S, C507S, C444S, C494S, C580S, C594S) that displayed reduced mature glycoform (**Fig. 3A&B**) and plasma membrane localization (**Fig. 4A&B**), also display greatly reduced or abolished prostaglandin uptake activity (**Fig. 5A**). The C450S variant displayed a ∼20% reduction in PGE_2_ transport activity compared to WT (**Fig. 5A**), which is consistent with this variant still displaying some plasma membrane localization (**Fig. 4A**).

**Figure 5.**
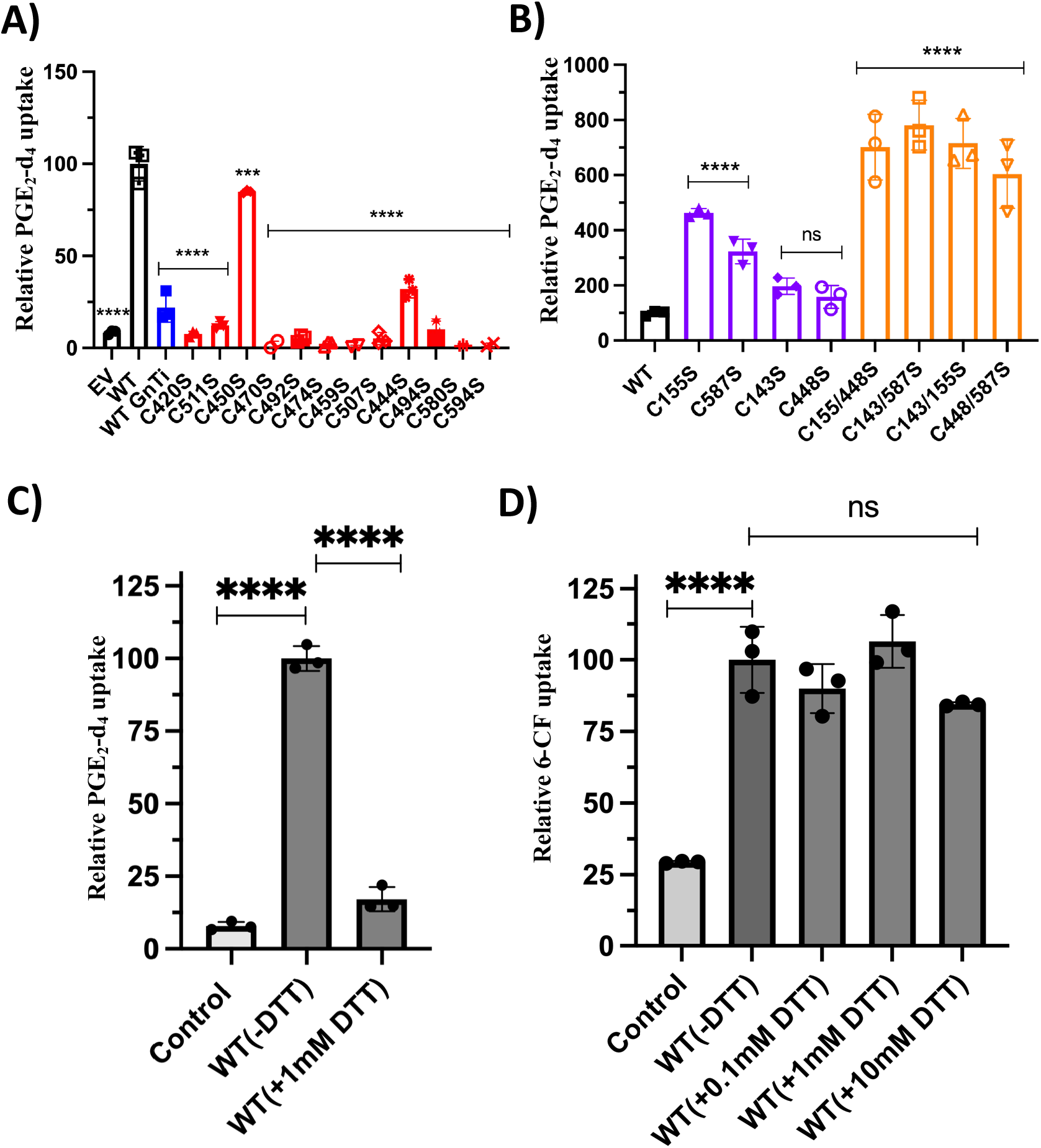
PGE_2_-d_4_ uptake activity of human PGT and cysteine variants. **A)** Reduced PGE_2_-d_4_ uptake activity of PGT single cysteine variants. The activity was normalized to WT PGT. Mock indicates the empty vector control. Error bars represent standard deviation of the mean (SD) across *n* = 3 triplicate measurements. Statistical significance was determined using ordinary one-way ANOVA, multiple comparison test and differences are depicted as \*\*\*\**P* ≤ 0.0001. **B)** Increased PGE_2_-d_4_ uptake activity of PGT single cysteine variants and their double cysteine variants. The activity was normalized to the surface expression level. **C)** Reduced PGE_2_-d_4_ uptake activity when PGT was treated with 1mM DTT. **D)** Reduced 6-CF uptake activity when PGT was treated with 0.1mM, 1mM and 10mM DTT.

In stark contrast, the four inter-loop cysteine variants (C155S, C587S, C143S, C448S) that displayed mature glycoform expression (**Fig. 3C&F**) and normal plasma membrane localization (**Fig. 4C**), possess PGE_2_ uptake activity that is increased by ∼2-5 fold over that of WT PGT (**Fig. 5B, purple columns**). Importantly, the uptake activity in **Fig. 5B** was normalized to the PGT plasma membrane expression level (**Fig. 4C**), and thus the increased uptake activity of these cysteine variants relative to WT transporter represents a true increase in uptake capacity rather than a difference in protein expression level or proportion of protein localized to the plasma membrane. These results suggest not only that inter-loop disulfide bonds between C143-C448 and C155-C587 are not essential for proper post-translational modification, membrane trafficking, or transport activity of PGT, but also that disulfide bonds between these cysteines may reduce maximal transport capacity for PGE_2_.

To further examine the role of inter-loop disulfide bonds between C155-C587 or C143-C448, we constructed double Cys-to-Ser variants (C155/448S, C143/587S, C143/155S, C448/587S) to explore all possible combinations of eliminating both potential disulfides. All four of these double cysteine variants are well-expressed with normal levels of mature glycoform (**Suppl. Fig. S8A**), and are localized to the plasma membrane similar to WT PGT (**Suppl. Fig. S8B**). Additionally, these double cysteine variants exhibit an even higher prostaglandin uptake activity than their individual cysteine variant counterparts, reaching as high as ∼7 fold higher uptake activity than WT PGT (**Fig. 5B, orange columns**). These results demonstrate that inter-loop disulfide bonds between C155-C587 and C143-C448 are not required for proper PGT maturation and trafficking, and may even diminish the maximal transport capacity of the transporter.

### Role of disulfide bonds after proper plasma membrane localization

Intra-loop disulfides in the ECD of PGT appear to be required for proper glycosylation **(Fig. 3D&E)** and trafficking **(Fig. 4A&B)** of the transporter, but it was not clear whether these disulfide bonds are required to maintain maximal transport capacity even after the transporter has been properly trafficked to the plasma membrane. To explore this possibility we performed further cellular uptake assays in the presence or absence of the reductant dithiothreitol (DTT). In these assays we monitored uptake of two alternative substrates, the fluorescent substrate 6-carboxyfluorescein (6-CF)^44^ or the native physiological substrate PGE_2_.

When HEK293 cells stably expressing WT PGT are subjected to treatment with 1 mM DTT, a significant decrease (∼80%) in total PGE_2_ uptake is observed (**Fig. 5C**). In contrast, titration of PGT-expressing HEK293 cells with increasing concentrations of DTT (0.1-10 mM) shows that the uptake activity of the exogenous substrate 6-CF remains the same as in WT cells without DTT treatment **(Fig. 5D)**. These results collectively demonstrate that disulfide bonds in the ECD are essential for maintaining maximal transport of some but not all substrates. A plausible explanation for these results is that disulfide bonds help maintain tertiary structure of the overall ECD, which is critical for transport of certain substrates such as PGE_2_.

## Discussion

PGT serves a dual role by mediating prostaglandin uptake from the extracellular environment, while also functioning as a core component of the Maxi-Cl channel to facilitate ATP and organic anion release upon interaction with the annexin A2-p11 complex. PGT is the only member of the OATP transporter family thus far that has been reported to exhibit such dual transporter-channel activity. The precise molecular mechanisms of prostaglandin import and ATP release mediated by PGT have remained poorly understood, partly due to the absence of structural investigations. Although recent cryo-EM structures of OATP1B1 and OATP1B3 have provided unprecedented insight into the structure and conformational states of OATP transporters, it remained unclear whether PGT with unique bimodal transporter/channel properties, shared the same structural topology as other OATP family members. Our cryo-EM structure presented here (**Fig. 1A**) demonstrates that PGT adopts an outward-facing state and shares the MFS fold characteristic of other OATP transporters (**Suppl. Fig. S5A&B**). Based on this structural similarity it is highly likely that PGT transports PGE_2_ and other small molecules across the membrane through conformational states like those observed previously for OATP1B1 and OATP1B3, however, further structural characterization of the PGT-A2-p11 complex is needed to clarify how PGT functions as a component of the Maxi-Cl channel.

We incubated PGT with the endogenous transport substrate PGE_2_ prior to grid preparation, but we were not able to observe this molecule in our cryo-EM maps. We suspect that bulky detergent molecules (**Fig. 1C and Suppl. Fig. S6**) may have obstructed the entry of PGE_2_ into the transport pocket. Although the exact binding mode of prostaglandins remains unresolved, our cryo-EM structure and molecular docking align with previous studies suggesting that the PGE_2_ binding pocket is near residues A526, A529, C530, and H533 in TM_10_ (**Suppl. Fig. S7A**)^45^. Mapping these residues onto our structure reveals their positioning along the central aqueous transport pathway, near the major ligand binding pocket identified in the OATP1B1-Bilirubin/Simeprevir bound structures^24^ (**Suppl. Fig. S7F-H**). Additionally, molecular docking identified PGE_2_ binding poses adjacent to these residues and near R561 (**Suppl. Fig. S7C&D**), which has previously been implicated as critical for substrate transport^46^. While a detailed molecular understanding of PGE_2_ binding and transport awaits further structural characterization, our cryo-EM structure and molecular docking provide a novel structural context upon which to interpret various earlier biochemical findings^45,46^.

In all currently available cryo-EM structures of OATPs the ECD displays consistently lower resolution and map quality than the transmembrane region, likely due to flexibility in these extracellular loops. Few investigations have characterized the ECD of OATP transporters, and the function of this domain across the entire family remains largely unknown. A notable feature of the ECD is the presence of 16 conserved cysteines that are proposed to form both intra- and inter-loop disulfide bonds. In PGT, missense mutations of several cysteines in the ECD have been shown to be causative for PHO and CEAS diseases^16,17^. Our results demonstrate that disulfide bonds in the ECD of human PGT are intricately linked to glycan modification and trafficking to the plasma membrane. Among the 8 putative disulfide bonds in the ECD, 6 are formed within a single loop (intra-loop), and are essential for proper maturation of glycan structures and subsequent trafficking to the plasma membrane **(Fig. 3A-B&4A-B)**. This result is consistent with previous findings where 10 cysteine mutations in the longest loop within the ECD of human OATP2B1 impede protein trafficking to the surface^23^. In contrast, the other two putative disulfide bonds in PGT (C155-C587 and C143-C448) are formed between loops (inter-loop) of the ECD and are not required for proper glycan maturation or plasma membrane localization **(Fig. 3C&4C)**. Our results thus demonstrate that intra-loop disulfide bonds in the ECD are essential for proper glycosylation and trafficking to the plasma membrane, while inter-loop disulfide bonds are dispensable for these processes.

Our examination of inter-loop disulfide bonds in the ECD (C143-C448 and C155-C587) showed an increase in the maximal PGE_2_ uptake when any of the four cysteines were mutated to serine. Breaking either inter-loop disulfide bond causes PGE_2_ uptake to increase ∼2-5 fold greater than WT (**Fig. 5B**). When both inter-loop disulfides were removed by a combination of double Cys-to-Ser mutations, we observed PGE_2_ uptake activity that was even further elevated to ∼6-8 fold above that observed for WT PGT (**Fig. 5B**). Thus, inter-loop disulfide bonds in the ECD of PGT appear to prevent maximal uptake of the substrate PGE_2_. This scenario raises an important question as to whether these two inter-loop disulfide bonds may play a regulatory role in controlling prostaglandin transport. Inflammation is intricately connected to cellular and tissue redox state^51,52^, and given that PGT serves as a key regulator of inflammatory signaling, it is plausible that redox-active enzymes such as thioredoxin or small molecules like glutathione^53^ that are secreted in response to inflammatory signals could play a role in reducing inter-loop disulfides. While it is conceivable that such a reduction could play a regulatory role by enhancing prostaglandin uptake to terminate signaling, further work in various cell types and redox environments will be required to reveal any such regulatory mechanism.

After PGT reaches the plasma membrane, disulfide bonds likely serve to maintain the overall tertiary structure of the ECD (largely the Kazal domain in loop-3). When DTT-induced disruption of disulfide bonds occurs, the uptake of endogenous PGE_2_ is significantly reduced **(Fig. 5C)**, while the uptake of the exogenous substrate 6-CF remains unaffected **(Fig. 5D)**. Thus, it appears that disulfide bonds are essential for maintaining an overall tertiary structure of the ECD that is critical for recognizing the endogenous substrate PGE_2_ before it is translocated to the transport pocket deep between the transmembrane helix bundles. In contrast, disulfide bonds and presumably the tertiary structure of the ECD are dispensable for maintaining transport of the surrogate substrate 6-CF. These results suggest that the ECD can play a critical role in recognizing some transported substrates, highlighting an underappreciated role of the ECD in mediating substrate selectivity of PGT and other OATP transporters.

Our study proposes a mechanistic model **(Fig. 6)** of the overall role of individual disulfide bonds within the ECD of PGT and likely other OATP transporters. First, six intra-loop disulfide bonds play an important role in the early folding and maturation of the transporter. These disulfides appear to be critical for proper maturation of glycan structures, which in turn influences trafficking of the mature transporter to the plasma membrane. Once the mature transporter is localized to the plasma membrane, these intra-loop disulfide bonds likely aid in maintaining correct folding of the ECD to facilitate prostaglandin uptake. In contrast, interloop disulfide bonds are dispensable for proper maturation and membrane trafficking, but their presence also reduces maximal PGE_2_ uptake capacity. The conserved nature of cysteine residues within the ECDs of human OATP transporters raises compelling questions about whether the folding and functional mechanisms we reveal here for OATP2A1 are a universal feature of this transporter family.

**Figure 6.**
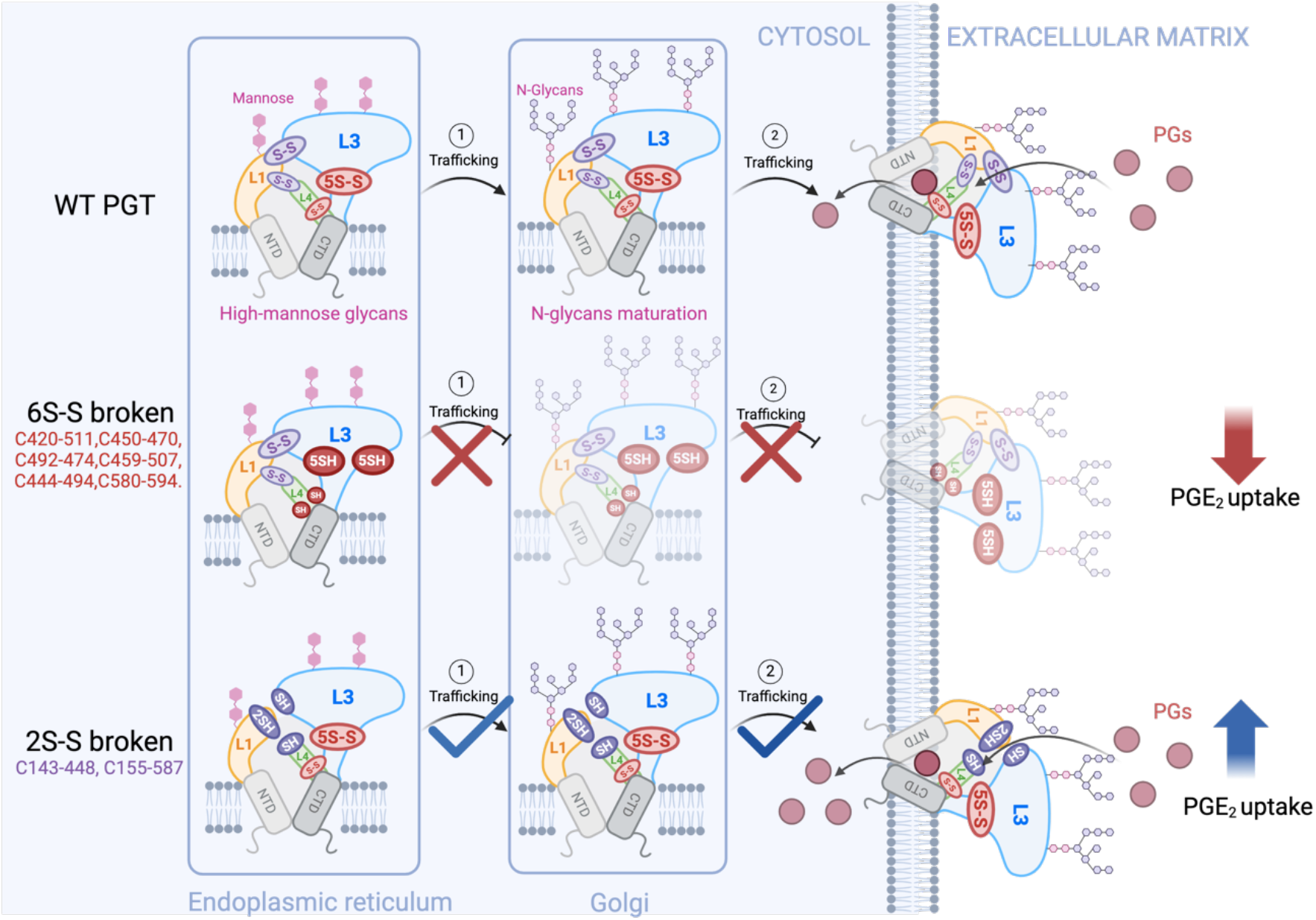
Model for ECD cysteines regulating PGT modification, trafficking, and function. Six intra-loop disulfide bonds (red circles) are critical for proper maturation of glycans and further influence the trafficking of mature PGT to the plasma membrane. Once the mature transporter is localized to the plasma membrane, these intra-loop disulfides maintain correct folding of the ECD to facilitate prostaglandin uptake. In contrast, the C143-C448 and C155-C587 inter-loop disulfide bonds (purple circles) appear to be dispensable for proper maturation of glycosylation and membrane trafficking of PGT, and rather seem to play a regulatory role by preventing maximal prostaglandin uptake.

## Materials and Methods

### Chemicals and Reagents

PGE_2_-d_4_ (CAS No. 34210-10-1) and PGE_2_-d_9_ (CAS No. 1356347-42-6) were purchased from Cayman Chemical (Ann Arbor, MI). High glucose Dulbecco’s Modified Eagle’s medium (DMEM), heat inactivated fetal bovine serum (FBS), and Penicillin/streptomycin were purchased from Gibco. HEK293-F cell line, GnTi cell line, Lipofectmine2000, EZ-Link™ Sulfo-NHS-SS-Biotin, *β*-Actin Monoclonal Antibody (BA3R)-HRP, and HA Tag Monoclonal Antibody (2-2.2.14)-HRP were purchased from ThermoFisher Scientific. NEB Q5 site-directed mutagenesis kit was purchased from NewEngland Biolabs. Strep-Tactin^XT^ resin was purchased from IBA Lifesciences. n-Dodecyl-β-D-Maltopyranoside (DDM) and Cholesteryl Hemisuccinate Tris Salt (CHS) were purchased from Anatrace. Digitonin was purchased from Millipore-Sigma.

### Cloning and Site-directed mutagenesis

The gene encoding human PGT was PCR-amplified from cDNA (TransOMIC Technologies, Alabama) and assembled into a pCDNA3.0 vector using NEB HiFi Assembly according to the manufacturer’s protocol. This process generated a plasmid containing human PGT with both N-terminal and C-terminal HA tags. Site-directed mutagenesis of human PGT was performed using a Q5 mutagenesis kit from NEB using standard manufacturer protocols. Plasmids encoding cysteine variants of human PGT were verified through Sanger sequencing at Azenta. Oligonucleotide sequences for cloning and mutagenesis are shown in **Suppl. Table S2**.

### Cell culture and transfection

HEK293-F cells were cultured in high glucose DMEM containing 10% FBS with penicillin (100 U/mL) and streptomycin (100 μg/mL) at 37°C in a humidified 7% CO_2_ atmosphere. Cells were seeded at a density of 0.5 × 10^6^ cells/well in 6-well plates and then transfected with plasmid constructs of wild-type (WT) PGT, cysteine variants, and pcDNA3.0 empty vector using lipofectamine 2000 reagent. Cells were further incubated for 48hrs to transiently express PGT and used for the following experiments including uptake assays and immunoblot analysis.

### Large scale expression and purification of PGT for structural studies

To express PGT for structural studies a HEK293F stable cell line was generated by geneticin (600ug/mL) selection of cells transfected with a pcDNA3.0 vector containing N-terminally twin-strep tagged PGT. The stable cell line was cultured in SMM 293-TII medium (Sino Biological) at 37 °C with 7% CO_2_ at 130 rpm in 3L non-baffled flasks on an orbital shaker. When the cell density reached approximately 2.0 × 10^6^ cells/mL, 4L of cells were harvested by centrifugation at 800× *g*. Cell pellets were resuspended with a dounce homogenizer in lysis buffer containing 50mM HEPES (pH 8), 300mM NaCl, 1 μg/mL pepstatin A, 1 μg/mL leupeptin, 1 μg/mL aprotinin, 0.6 mM benzamidine, 5kU benzonase nuclease, and solubilized by stirring for 1 h at 4 °C with 1% (w/v) DDM and 0.2% (w/v) CHS. After solubilization the cell lysate was centrifuged at 100,000× *g* for 1 h and the supernatant was loaded by gravity flow onto 1mL of Strep-tactin^XT^ resin. The resin was sequentially washed with 10CV of buffer A (50 mM HEPES (pH 8), 300 mM NaCl) with 0.05% (w/v) DDM and 0.01% (w/v) CHS, followed by 5CV of buffer A with 0.06% (w/v) digitonin, before being eluted with buffer A containing 0.06% (w/v) digitonin and 50mM biotin. The protein sample after elution was concentrated in a 100kDa MWCO concentrator and further purified by size-exclusion chromatography on a Superose 6 10/300 GL column in 25mM HEPES, 150mM NaCl, 0.06% digitonin. Peak fractions were analyzed by SDS-PAGE and concentrated to ∼3.8 mg/mL for cryo-EM analysis.

### Cryo-EM imaging and data processing

Grids for cryo-EM imaging were prepared on a Vitrobot Mark IV by applying 3 μL of purified PGT to Quantifoil R1.2/1.3 200 mesh grids that had been glow-discharged for 45 s at 15 mA in a Pelco EasyGlow. Grids were blotted for 3-5 s at 4 °C, 100% humidity before being plunge frozen in liquid ethane cooled by liquid nitrogen. Data collection was performed on a Talos Arctica equipped with a Selectris energy filter and Falcon 4i Direct Electron Detector. Movies were collected in counting mode with a pixel size of 0.886Å, a total dose of 44.48 electrons/Å^2^ and a defocus range from - 0.5μm to -2μm. Movies were corrected for beam-induced motion by performing patch motion correction followed by patch CTF estimation in cryoSPARC^32^. Obviously poor micrographs and those with a CTF fit worse than 5Å were excluded from further analysis. Blob picking was used to generate initial 2D templates for template-based particle picking in cryoSPARC. Several rounds of 2D classification were performed to remove poor particles, and initial models for 3D classification were constructed using *ab initio* reconstruction in cryoSPARC using frequencies in the range of 12-8Å. Following heterogeneous classification and non-uniform refinement^33^ in cryoSPARC, selected particles were transferred to RELION 5.0^34^ for further 3D classification and refinement with Blush regularization^34^. Resolutions of all maps were calculated using the gold-standard FSC at a cutoff of 0.143.

The AlphaFold^35^ model of outward-facing PGT was initially rigid-body docked into the WT cryo-EM map using UCSF Chimera^36^. The model was manually adjusted in COOT^37^ to properly fit the cryo-EM density and refined in real-space using phenix.real_space_refine in the PHENIX software suite^38^. Iterative rounds of real-space refinement in PHENIX and manual model adjustment in COOT were performed to optimize the overall model geometry and fit to the experimental cryo-EM map. All models were assessed for appropriate stereochemical properties and fit to the experimental cryo-EM map using MolProbity^39^ as implemented in PHENIX (**Suppl. Table S1**). Figures of cryo-EM maps and atomic models were created using UCSF Chimera^36^ or UCSF ChimeraX^40^. BioRender was used to make images throughout the manuscript and supplemental material.

### Ligand docking

Molecular docking was performed using AutoDock Vina^41^. A structural model of deprotonated PGE_2_ was obtained from PubChem (CID 11865423) and docked against the structure of PGT as determined from cryo-EM. Nonpolar hydrogens were added to the protein model prior to ligand docking. Flexible docking was performed by defining a set of six flexible residues that had previously been shown to be involved in ligand binding (Arg561, Lys614, Ser526, Ala529, Cys530 and His533). The grid box was prepared in AutodockTools^42^ with a box size of ∼40Å^3^ centered around the central transport pore of PGT. An exhaustiveness of 64 was used to ensure extensive pose sampling, and the top three poses were selected.

### Cell surface biotinylation and immunoblot analysis

HEK293-F cells were cultured in a 6-well plate and transiently transfected with pcDNA3.0 vectors containing WT PGT or site-directed variants. At 48 h post-transfection the cells were washed twice with 1 mL of ice-cold uptake buffer (25mM HEPES, 125mM NaCl, 4.8mM KCl, 5.6mM Glucose, 1.2mM CaCl_2_, 1.2mM KH_2_PO4, 1.2mM MgSO_4_, pH 7.4) and then treated for 1 h at room temperature with 1 mL of cell impermeable EZ-Link™ Sulfo-NHS-SS-Biotin (0.5mg/ml in uptake buffer). After this, cells were washed three times with 1 mL of ice-cold uptake buffer containing 50 mM glycine and incubated for 10 min at 4 °C. Cells were subsequently lysed with 300 μL of lysis buffer (25mM HEPES, 150 mM NaCl, 1% DDM, 0.2% CHS, pH 7.4, containing protease inhibitors and nuclease) for 1 h at 4 °C with shaking. Lysates were centrifuged for 10 minutes at 17,000×g, and the supernatants were incubated with 50 μL of Strep-Tactin beads for 1 hour at 4°C under constant agitation. The beads were then centrifuged at 100× g for 30s, washed for three times with wash buffer (25mM HEPES, 150mM NaCl, 0.05% DDM, 0.01% CHS), and incubated with 100 μL of wash buffer containing 100 mM DTT at room temperature for 30 min to recover biotinylated surface proteins. Samples were separated using SDS-PAGE followed by western blot analysis. Briefly, SDS-page separated protein were transferred to a nitrocellulose membrane, blocked in 5% dry milk, and probed with an anti-HA-HRP antibody (1: 3000 dilution). WT PGT and cysteine variants were detected using a ECL™ prime western blotting detection kit and chemiluminescent detection on BioRad imaging system.

### Ligand uptake assays

PGT-mediated transport activity in HEK293-F cells was analyzed using PGE_2_-d_4_ as a substrate to distinguish it from endogenous PGE_2_ as previously described^43^. Cells were incubated in uptake buffer containing 1uM PGE_2_-d_4_ for 5 min at 37°C. Cells were subsequently washed three times with ice-cold uptake buffer, before being lysed with 70% acetonitrile containing PGE_2_-d_9_ as an internal standard. The samples were centrifuged and the supernatant was dried under speed vacuum. The residue was reconstituted with 0.1% formic acid/acetonitrile (3:1, v/v) and separated on a Waters Acquity HPLC system equipped with a Waters Acquity UPLC BEH C18 (1.7um 2.1*100mm) analytical column. Mobile phase A consists of 0.1% acetic acid, and mobile phase B is composed of 16% methanol, 84% acetonitrile, and 0.1% acetic acid. The gradient begins at a 65:35 ratio of A:B and transitions to a 1:99 ratio of A:B. The flow rate of the mobile phase was 0.3 mL/min, and the injection volume was set as 5 µL. Samples eluting from the column were analyzed on a Waters Xevo TQ-XS triple quadrupole mass spectrometer using electrospray ionization in negative ion mode. Mass transitions were monitored at m/z 355.20/275.20 for PGE_2_-d_4_ and 360.20/280.20 for PGE_2_-d_9_. MassLynx XS software was used for data analysis.

The 6-CF uptake assay^44^ was conducted in an uptake buffer at pH 6.0 (25 mM MES, 125 mM NaCl, 4.8 mM KCl, 5.6 mM glucose, 1.2 mM CaCl_2_, 1.2 mM KH_2_PO_4_, 1.2 mM MgSO_4_). Cells were incubated with 100uM 6-CF in uptake buffer for 5 minutes, followed by two washes with PBS. The cell pellet was then resuspended in PBS, and the accumulation of 6-CF was measured by fluorescence using a microplate reader (SpectraMax iD5) with excitation at 485 nm and emission at 535 nm. For DTT treatment, cells were incubated with varying concentrations of DTT in PBS for 20 minutes, followed by washing with PBS before performing the uptake assay.

## Data availability

Atomic coordinates and associated electron microscopy maps for the human PGT structure reported in this publication have been deposited in the Protein Data Bank (PDB) and Electron Microscopy Data Bank (EMDB) under the following accession numbers: (PDB: 9MGK and EMDB: 48261).

## Acknowledgments

Research reported in this publication was supported by departmental startup funds and the James K. Billman Jr. MD Endowed Research Professorship to B.J.O. The content is solely the responsibility of the authors. We would like to thank Dr. Sundharraman Subramanian for help with electron microscope operation at the RTSF Cryo-Electron Microscopy Core at Michigan State University, and Dr. Tony Schilmiller for assistance with mass-spectrometry data collection and analysis at the RTSF Mass-Spectrometry and Metabolomics core. We would also like to thank all members of the Orlando laboratory for detailed discussions and analysis of the manuscript.

