## Supplemental Information for "Structure and Post-Translational Modification of the Prostaglandin Transporter"

**This PDF file includes:**

Figures S1 to S8
Tables S1 and S2

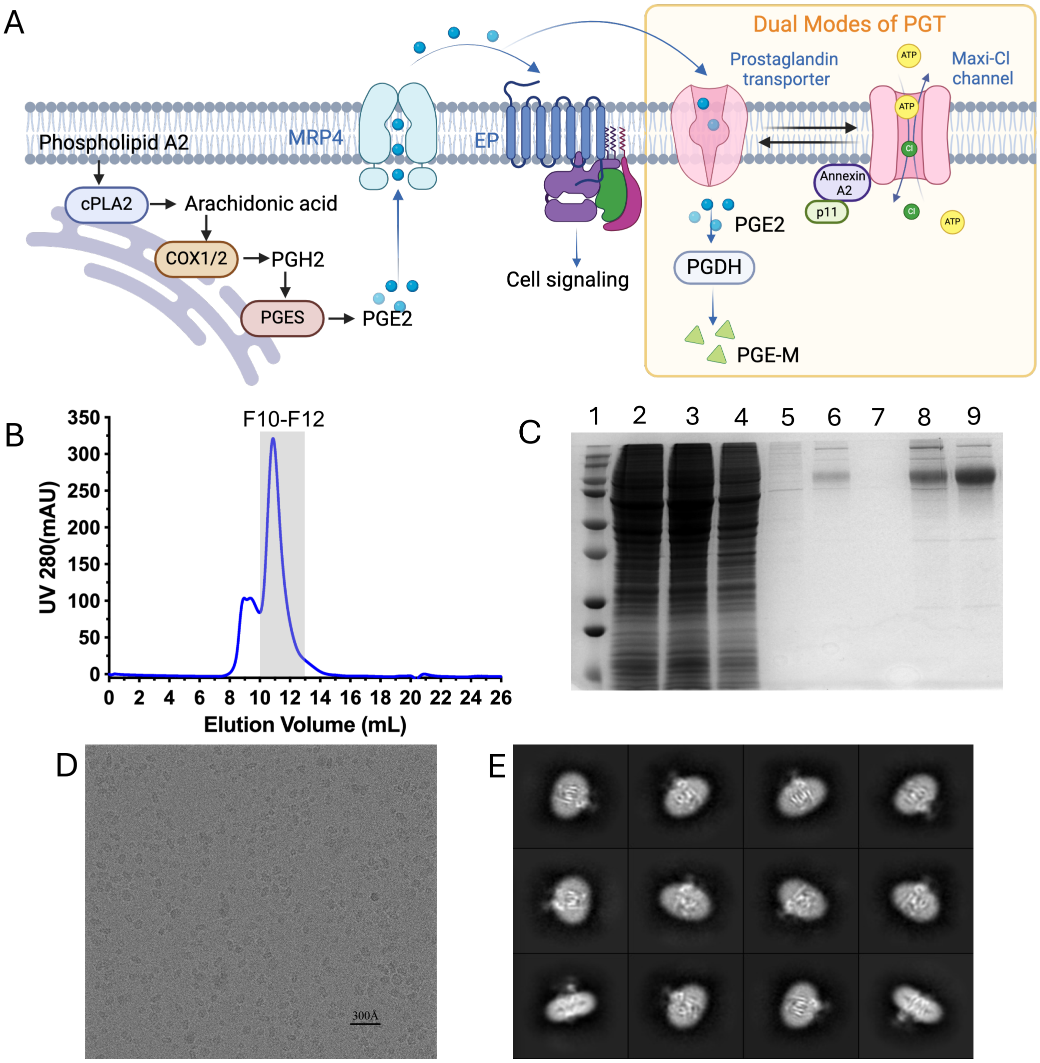


**Fig. S1. Purification and biochemical characterization of PGT**

**A)** Schematic of prostaglandin biosynthesis and metabolism. Prostaglandin precursors are synthesized in the cytosol, transported outside the cell to trigger cellular responses, and then transported back into the cytosol for degradation. cPLA2: cytosolic phospholipase A2. COX1/2: cyclooxygenase enzyme 1/2; PGES: prostaglandin E synthase; EP: Prostaglandin GPCR receptors; PGDH: prostaglandin dehydrogenase. **B)** Gel-filtration profiles of detergent solubilized apo PGT. **C)** SDS-PAGE analysis of purified PGT. Lane 1: protein ladder; Lane 2: supernatant; Lane 3: Flow through; Lane 4 and 5: Wash; Lane 6: Elution from Strep-Tactin column; Lane 7: Concentrator flow through; Lane 8: Concentrated elution before gel-filtration; Lane 9: Gel-filtration F10-F12. **D)** Electron micrograph of detergent solubilized PGT showing a uniform particle distribution in thin ice. The black scale bar demonstrates a size of 300Å. Approximately ~9000 similar micrographs were obtained. **E)** Representative 2D classification of PGT.

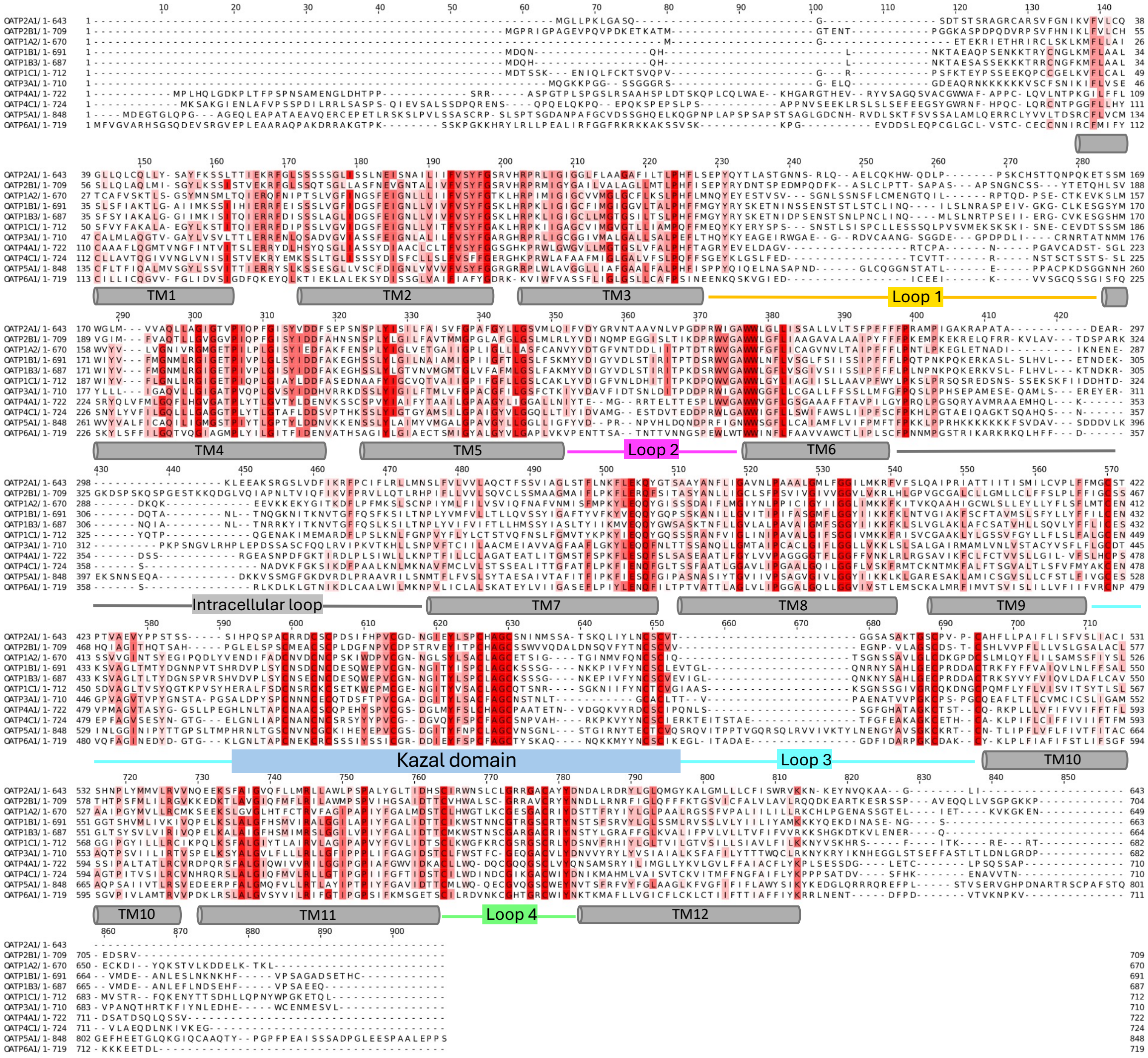

**Fig. S2. Sequence alignment of 11 human OATPs**

Clustal Omega sequence alignment of OATP2A1 (PGT)(Homo sapiens OATP2A1, ID: Q92959), OATP2B1 (Homo sapiens OATP2B1, ID: O94956), OATP1A2 (Homo sapiens OATP2B1, ID: P46721), OATP1B1 (Homo sapiens OATP1B1, ID: Q9Y6L6), OATP1B3 (Homo sapiens OATP1B3, ID: Q9NPD5), OATP1C1(Homo sapiens OATP1C1, ID: Q9NYB5), OATP3A1(Homo sapiens OATP3A1, ID: Q9UIG8), OATP4A1(Homo sapiens OATP4A1, ID: Q96BD0), OATP4C1(Homo sapiens OATP4C1, ID: Q6ZQN7), OATP5A1(Homo sapiens OATP5A1, ID: Q9H2Y9) and OATP6A1(Homo sapiens OATP6A1, ID: Q86UG4) sequences. The motifs and secondary structure elements are depicted. TM: transmembrane helix.

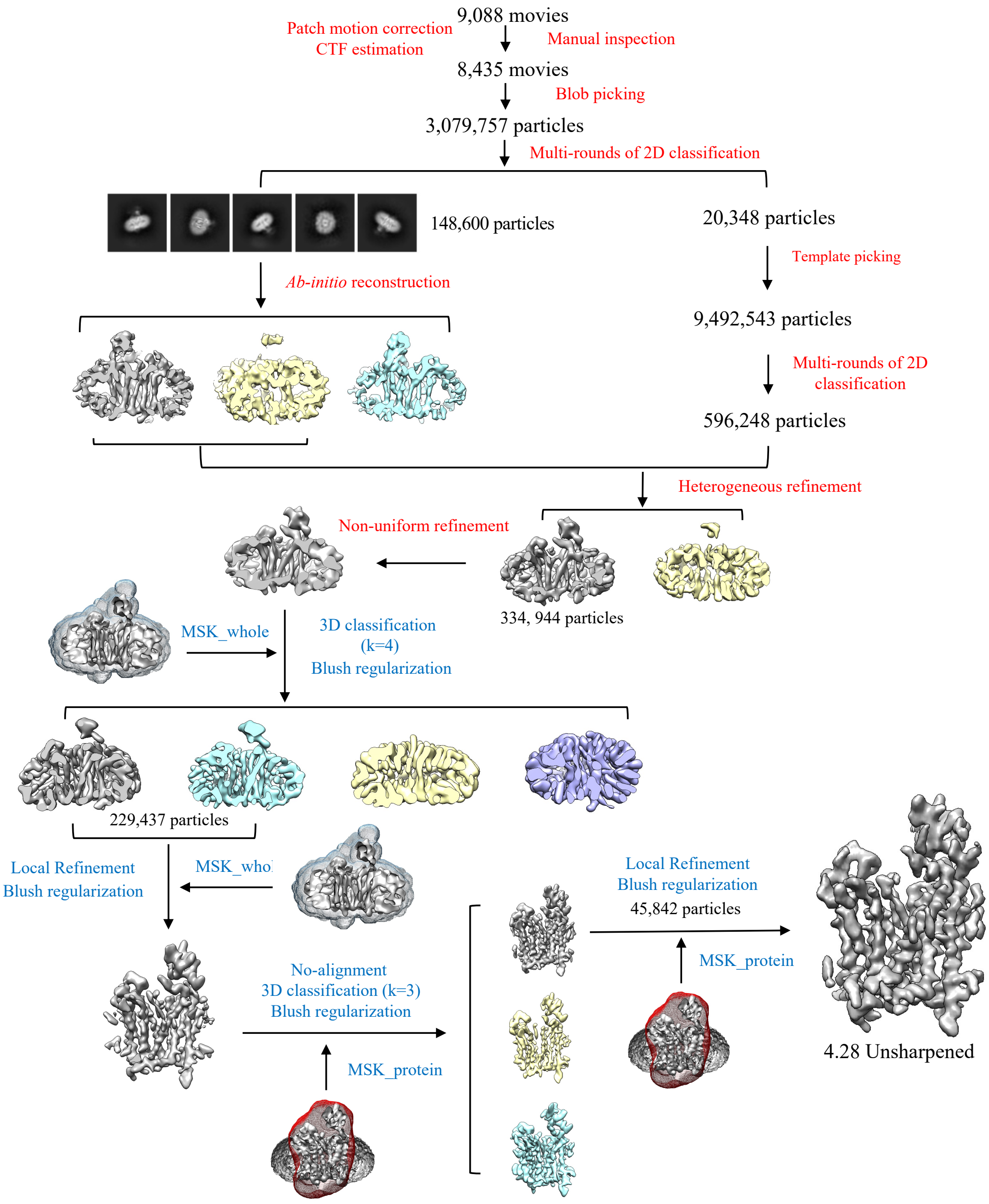


**Fig. S3. Cryo-EM data processing workflow**
Steps shown in red were performed in CryoSPARC and steps shown in blue were performed in Relion. Particle numbers in each step are written in black.

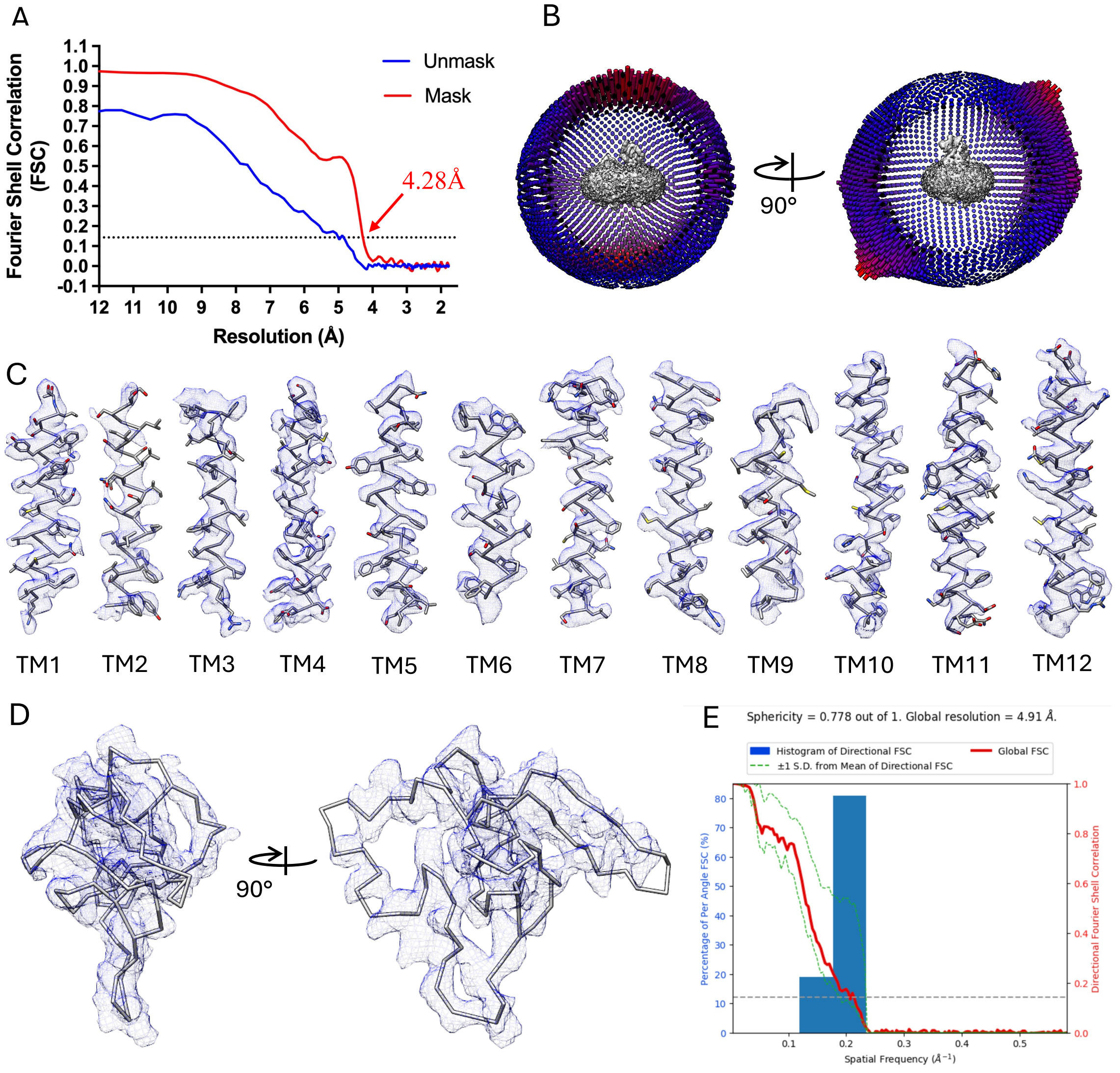


**Fig. S4. Cryo-EM data analysis for PGT**

**A)** FSC curve for the final 3D reconstruction of PGT. Blue curve shows the FSC between unmasked half-maps, red curve shows FSC between the same halfmaps after applying a tight mask in CryoSPARC. Gold standard FSC=0.143 is indicated with a dotted black line. **B)** Angular distribution of particles in the final reconstruction of PGT. **C)** Coulomb potential map for each individual transmembrane helix of PGT. **D)** Coulomb potential map for extracellular loop-3 Kazal domain of PGT. **E)** 3D FSC as calculated from the 3DFSC web server.

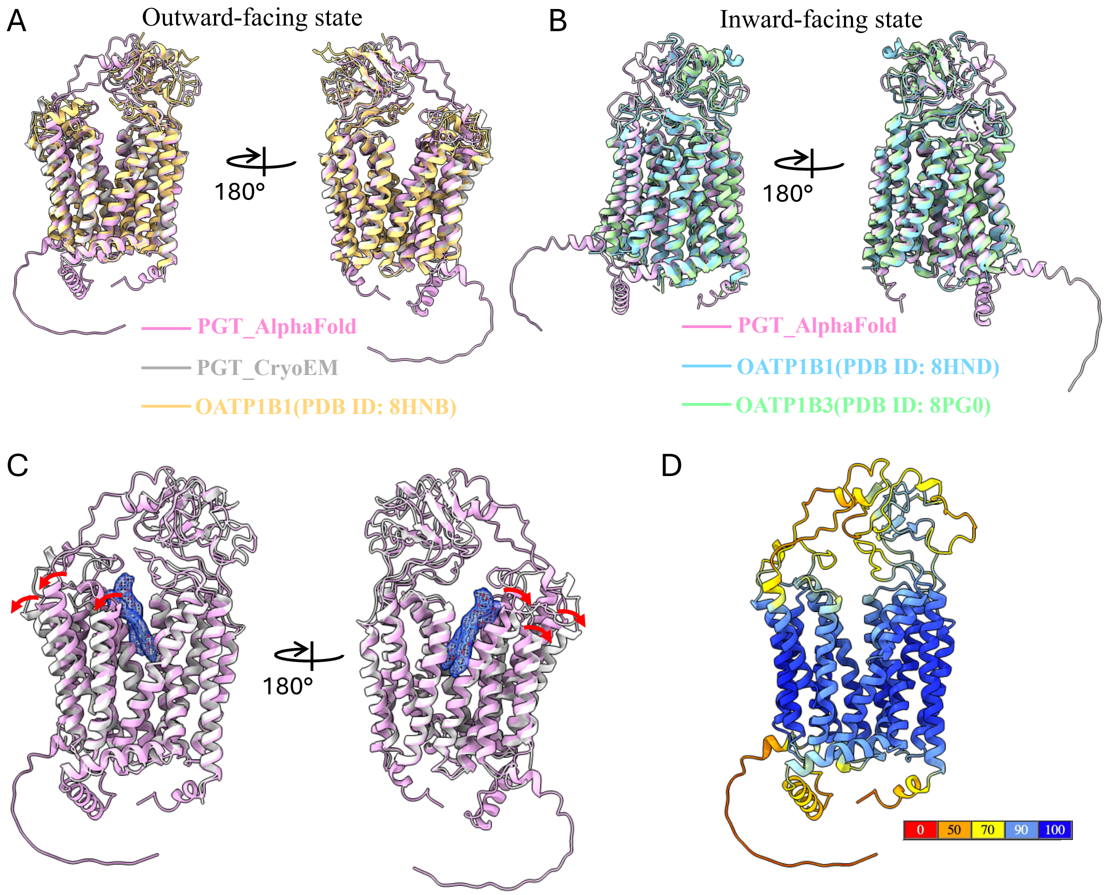


**Fig. S5. AlphaFold predicted PGT conformations and potential substrate binding pocket**

**A)** Structural alignment of cryo-EM-determined PGT (grey), AlphaFold-predicted PGT (pink), and cryo-EM-determined OATP1B1 (orange) in an outward-facing state. **B)** Structural alignment of AlphaFold-predicted PGT (pink), cryo-EM-determined OATP1B1 (blue) and cryo-EM-determined OATP1B3 (green) in an inward-facing state. **C)** Detergent binding causes the N-terminal half of the CryoEM-PGT (light grey) open wider than AlphaFold PGT (Pink). **D)** AlphaFold predicted PGT model, colored by pLDDT score.

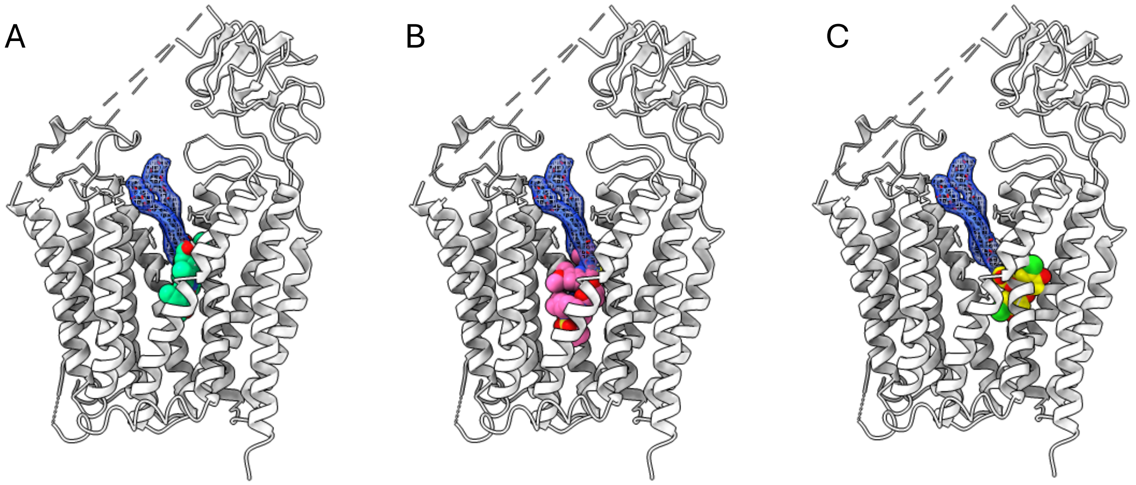


**Fig. S6. Alignment of PGT with OATP1B3 bound to small-molecules**

**A)** Bilirubin (green spheres) bound OATP1B1 (PDB ID: 8HNC) overlay with PGT. **B)** Semiprevir (pink spheres) bound OATP1B1 (PDB ID: 8HNH) overlay with PGT. **C)** Dichlorofluorescein (yellow spheres) bound OATP1B1 (PDB ID: 8K6L) overlay with PGT. The cryoEM derived PGT model is shown in grey cartoon in all panels, with the digitonin molecules observed in the PGT cryo-EM map shown in blue mesh.

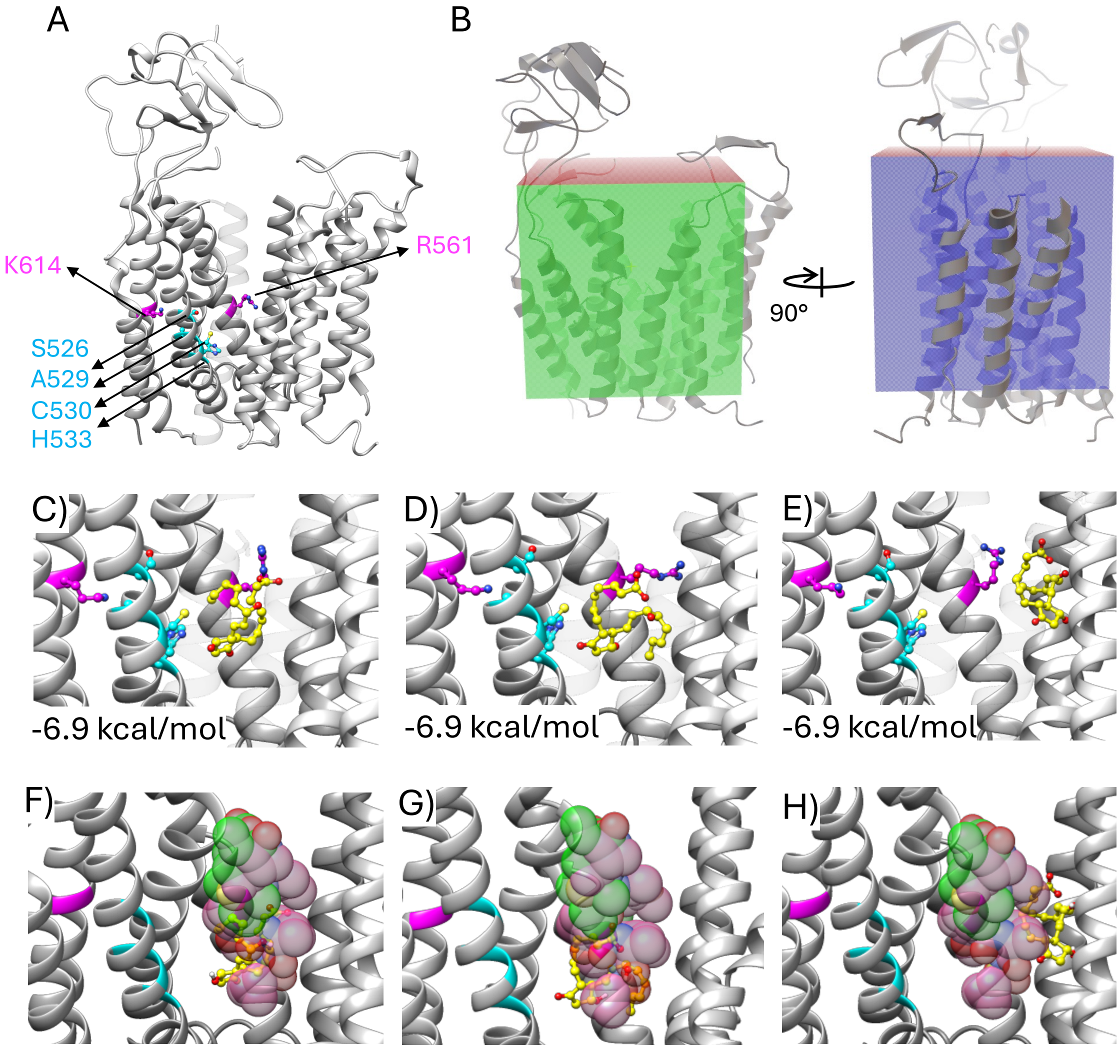


**Fig. S7. Molecular docking of $PGE_2$ to PGT**

**A)** Previous biochemical characterization identified R561 and K614 (magenta) as critical determinants for binding anionic substrates, and residues S526, A529, C530, and H533 (cyan) as lining the $PGE_2$ binding pocket. **B)** Diagram showing the ~40Å$^3$ gridbox designating the area for docking of $PGE_2$. **C-E)** Top 3 $PGE_2$ poses generated from docking of $PGE_2$ to PGT. **F-H)** $PGE_2$ poses from molecular docking overlaid with Bilirubin and Semiprevir bound OATP1B1. Yellow: $PGE_2$; Green sphere: Bilirubin; Pink sphere: Semiprevir.

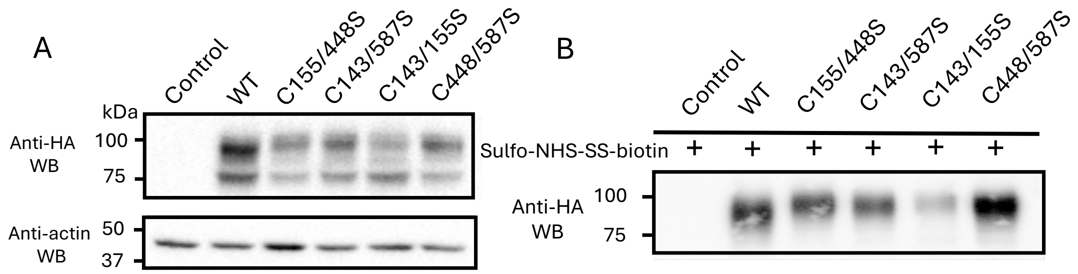


**Fig. S8. Total expression level and surface expression of PGT double cysteine variants**
**A)** Immunoblot of whole cell expression level of WT PGT and double cysteine variants. **B)** Immunoblot of surface expression level of WT PGT and its double cysteine variants.

**Table S1. Cryo-EM data collection, refinement, and validation statistics**

| | Human PGT (PDB: 9MGK) (EMDB: 48261) |
|---|---|
| **Data collection and processing** | |
| Magnification | 130,000 |
| Voltage (kV) | 200 |
| Electron exposure (e-/Å) | 44.48 |
| Defocus range ($\mu$m) | -0.5 ~ -2.0 |
| Pixel size (Å) | 0.886 |
| Symmetry imposed | C1 |
| Number of micrographs (#) | 9088 |
| Map resolution (Å) | 4.3 |
| FSC threshold | 0.143 |
| **Refinement** | |
| Model resolution | 4.4 |
| FSC threshold | 0.5 |
| Map sharpening *B* factor (Å$^2$) | -100 |
| Model composition | |
| Non-hydrogen atoms | 4135 |
| Protein residues | 528 |
| Ligands | 2 |
| Mean *B* factors (Å$^2$) | |
| Protein | 66.44 |
| Ligand | 43.93 |
| R.m.s.deviations | |
| Bond lengths (Å$^2$) | 0.003 |
| Bond angles (°) | 0.716 |
| Validation | |
| MolProbity score | 2.08 |
| Clashscore | 13.69 |
| Poor rotamers (%) | 0.00 |
| Ramachandran plot | |
| Favored (%) | 93.22 |
| Allowed (%) | 6.78 |
| Disallowed (%) | 0.00 |

**Table S2. Oligonucleotide primers used to generate human PGT cysteine variants**

| Variants | Forward Primers | Reverse Primers |
|---|---|---|
| C420S | CTTTGTTCTTCATGGGAtccTCCACCC | GAACACAAAGGATCATGGAGATGGTGATGATG |
| C511S | CCCTGTCCCCtctGCCCACTTCC | CACGATCCTGTCTTTGCTGAAGC |
| C450S | GGACTGCTCGtccCCAGATTCTATCTTC | CTGCGGCAGGCAGGAGAC |
| C470S | CCTCTCCCCTtccCATGCCGGCT | TACTCGATTCCATTGTCTCCACAGACC |
| C492S | CTATTTGAACtccAGCTGTGTGACCG | ATCAGTTGCTTGGAGGTTGCAGAGCTC |
| C474S | CCATGCCGGCtccAGCAACATCA | CAAGGGGAGAGGTACTCGATTC |
| C459S | CACCCGGTCtctGGAGACAATGG | GAAGATAGAATCTGGGCACGAGCAGTC |
| C507S | GATCGtccCCTGTCCCCTGTG | CTGTCTTTGCTGAAGCGGATCCCC |
| C444S | GTCTCCTGCCtccCGCAGGG | TGCGGATGTATAGAACTTGATGTGCTAGGG |
| C494S | GAACTGCAGCtctGTGACCGGG | AAATAGATCAGTTGCTTGGAGGTTGCAGAGC |
| C580S | CCATTGACCACTCCtccATCCGG | TGAGGCCATAGAGGGCTGGAGATG |
| C594S | GCGAGGGGCCtccGCCTACTATG | CTCCCCAAGCACAGCGAG |
| C143S | GGCCGAGCTCtccCAGAAG | TGCAAGCGGCTGTTGTTCCCAGTG |
| C448S | CCGCAGGGACtccTCGTGCCCAG | CAGGCAGGAGACTGCGGATG |
| C155S | CCCAGTAAGtccCACAGCACC | AGGCAGGTCCTGCCAATGCTTCT |
| C587S | GGAACTCGCTGtccTTGGGGAG | ACCGGATGCAGGAGTGGTCAATGG |
